# Graph Theoretical Network Structures Underlie Age-Related Differences in the Functional Connectome During Rest and Music Listening

**DOI:** 10.1101/2023.08.08.552499

**Authors:** Alexander Belden, Psyche Loui

## Abstract

Aging is associated with decreased functional connectivity within the default mode network, as well as auditory and reward systems which are involved in music listening. Understanding how music listening affects network organization of the aging brain, both globally and specific to the brain networks, will have implications for designing lifestyle interventions that tap into distinct networks in the brain. Here we apply graph-theory metrics of modularity, global efficiency, clustering coefficients, degrees, and betweenness centrality to compare younger and older adults (YA/OA, N=24 per group) in fMRI connectivity during rest and a music listening task. Results show a less modular but more globally efficient connectome in OAs, especially during music listening, resulting in main effects of group and task, as well as group-by-task interactions. ROI analyses indicated that the posterior cingulate is more centrally located than the medial prefrontal cortex in OAs. Overall, reduced modularity and increased global efficiency with age is in keeping with previously-observed functional reorganizations, and interaction effects show that age-related differences in baseline network organization are reflected in, potentially magnified by, music listening.

## Introduction

Change in metabolic activity across multiple brain networks is a consistent pattern observed in the study of aging. This is primarily characterized by decreased connectivity between regions of similar known function, and increased connectivity between functionally distinct regions (Grady et al., 2016; Sala-Llonch et al., 2015; Tomasi & Volkow, 2012). For example, aging has been associated with functional dysregulation of the default mode network (DMN) (Buckner et al., 2005; Hafkemeijer et al., 2012; Sheline et al., 2010), auditory network (Belden et al., 2023; Onoda et al., 2012), and reward network (Erixon-Lindroth et al., 2005; Li & Rieckmann, 2014), among others. In particular, dysregulation of the DMN, specifically the medial prefrontal cortex (mPFC), is strongly associated with age-related cognitive decline (Buckner et al., 2005; Hafkemeijer et al., 2012; Sheline et al., 2010). Ventral portions of the mPFC are also involved in reward learning (Knutson et al., 2003; O’Doherty et al., 2003), a process involving the release of dopamine during cognitively demanding tasks (Aalto et al., 2005; Badgaiyan et al., 2007; Koepp et al., 1998; Monchi et al., 2006).

While dopaminergic responses to task demands are preserved throughout the lifespan, older adults (OAs) show decreased dopamine release in cognitively demanding contexts relative to younger adults (YAs) (Bäckman et al., 2010; Eppinger et al., 2012; Samanez-Larkin et al., 2014; Yee et al., 2019). Dopamine release also shows a direct relationship with task performance (MacDonald et al., 2009) and Striatal BOLD response (Schott et al., 2008). Reward learning is a critical process that involves the formation of cognitive representations of predictions and prediction errors (Schultz et al., 1997), as well as affective, value-based representations of rewarding stimuli (Mather, 2016). Cognitive changes in reward learning associated with aging include a decrease in the ability to form prediction errors (Chowdhury et al., 2013; Samanez-Larkin et al., 2014), as well as a general deficit in working memory capacity and maintenance (Klostermann et al., 2012; Landau et al., 2009). Still, some report affective components of reward learning to be relatively preserved in aging populations (Geddes et al., 2018; Samanez-Larkin et al., 2014).

Considering these mixed reports, it would be of great interest to observe how rewarding experiences may stimulate or reorganize the functional connectome, and how such stimulation or reorganization may change with age. Music offers a rich and rewarding set of stimuli that engages multiple brain systems (Ai et al., 2022; Chen et al., 2022; Global Council on Brain Health, 2020; Zhang et al., 2023). Music listening, perhaps the most frequent form of music experience in daily life, can lead to increased functional connectivity throughout the brain (Karmonik et al., 2016; Ueda et al., 2013), including auditory-motor (Palomar-García et al., 2016) and auditory-reward connectivity (Alluri et al., 2017; Belden et al., 2023; Quinci et al., 2022). Sensitivity to the rewards of music listening also varies at the individual level: Higher musical reward, as measured through the Barcelona Music Reward Questionnaire (BMRQ) (Mas-Herrero et al., 2014), has been associated with increased connectivity between auditory and reward systems (Martínez-Molina et al., 2016; Sachs et al., 2016; Salimpoor et al., 2013), and thus musical reward may be a strong explanatory factor for individual differences in functional responses to music.

Music is also thought to be used as a tool for mood regulation in OA populations (Laukka, 2007), and receptive music-based interventions (MBIs) centered around preferred music listening have been shown to improve depression and anxiety in OAs with or at risk for Alzheimer’s Disease (AD) and other forms of dementia (Fang et al., 2017; Gaviola et al., 2020; Leggieri et al., 2019). However, effect sizes for improvement of cognitive function have been small, due in part to a large variance of outcomes (Ueda et al., 2013). By better characterizing the ways in which the brain responds to music, and in particular characterizing ways in which these responses change in older adulthood, we can better understand the systems that might be driving the positive effects of these interventions, which may in turn better inform the development of future MBIs (Chen et al., 2022).

Previous work has shown that when compared to OAs, YAs have higher within-network connectivity in auditory and reward systems during rest and music listening (Belden et al., 2023). This included a pattern of higher connectivity between auditory network and mPFC in YAs that was specific to music listening. Additionally, OAs showed global patterns of decreased anticorrelations to out-of-network regions, including a consistent pattern of higher connectivity to posterior cingulate cortex (PCC) in OAs. However, this study was focused on connectivity from particular systems of interest (auditory and reward networks), and did not specifically measure differences in connectome-wide patterns of connectivity. nor did it target other well-defined networks, such as the DMN. Here, we aim to characterize age and task-related differences across the functional connectome and specifically compare the DMN hubs of mPFC and PCC through the use of graph-theoretical analysis.

Graph-theoretical analyses of the functional connectome can quantify connectivity patterns spanning the entire brain, as well as the roles that specific regions of interest may play within the connectome’s topological structure (Farahani et al., 2019; Rubinov & Sporns, 2010). One graph-theoretical property of brain networks is modularity, which describes a graph’s tendency to assort into smaller subnetworks. Modularity has been shown to decrease with age (Geerligs et al., 2015; Onoda & Yamaguchi, 2013). Global efficiency, which is the average of inverse distances between all pairs of nodes, has not been shown to differ across age groups (Geerligs et al., 2015). Lower modularity in OAs could be partially explained by previously reported decreases in within-network and increases in out-of-network connectivity associated with age, although up until this point no work has determined how music listening may modulate these age-related differences.

One possibility is that music listening, through its engagement of numerous brain systems, could help restore the normal connectivity patterns of these systems, leading to an increase in modularity in OAs. Alternatively, music’s simultaneous engagement of multiple systems could result in less distinct boundaries between networks, potentially leading to decreased modularity. However, we would expect such an effect to have a stronger impact on those displaying fewer out-of-network connections at rest, in this case YAs, similarly resulting in a reduction of age-related differences in modularity. Given observed dysregulation of the mPFC associated with age (Buckner et al., 2005; Hafkemeijer et al., 2012; Sheline et al., 2010), as well as increased recruitment of the PCC by various systems with age (Belden et al., 2023), we could also expect that the connectivity profile of these regions could be notably different between age groups.

Here we use local graph theoretical metrics, namely degrees, clustering coefficients (CC), and betweenness centrality (BC) (the size of the region’s neighborhood, how tightly connected the neighborhood is, and how centrally located the region is within the overall graph, respectively) to quantify these regions’ contributions to the connectome’s overall graphical structure. A node is considered to be a hub if it has notably more connections than other nodes within a graph, showing a higher degree and betweenness centrality and a lower clustering coefficient when compared to other nodes. Here, we expect that the mPFC may have a more hublike function in YAs, while OAs may instead show more hublike qualities in PCC.

In this secondary analysis, we compare graph theoretical network structure of fMRI data collected during rest and music listening in a sample of young and OAs (OSF Preregistration: https://osf.io/25k9t). Our hypotheses were threefold:

Hypothesis 1: Considering whole-brain measures of modularity, in keeping with previous reports (Geerligs et al., 2015; Onoda & Yamaguchi, 2013), we predicted that YAs would show higher modularity than OAs. We also expected that modularity in OAs –– but not YAs –– would increase during music listening, resulting in an age-by-task interaction effect on modularity.
Hypothesis 2: Specifically within the DMN, we predicted that age-related differences in connectivity of mPFC and PCC would be reflected in measures of betweenness centrality and clustering coefficients. We expected that a more hublike structure of mPFC in YAs would lead to increased BC and decreased CC relative to OAs, and that the inverse would be true for PCC.
Hypothesis 3: Finally, we predicted auditory and reward networks to have higher global efficiency in participants with higher self-reported musical reward. We expected this effect to be present during both rest and music listening, and to be particularly robust when listening to well-liked and self-selected music.

## Methods

### Participants

Hypotheses 1 and 2 were tested by comparing samples of 24 OAs (20 female, 3 male, 1 non-binary) aged 54-89 (M =66.67, SD=7.79) and 24 YAs (11 female, 13 male) aged 18-23 (M=18.58, SD=1.21) as described in Belden et al (2023). YA participants were recruited through the Northeastern University student population and met the following inclusion criteria: 1) were 18 years of age or older, 2) had normal hearing, and 3) and passed an MRI screening. Twenty-eight OAs were recruited as part of a longitudinal study (Quinci et al., 2022) and met inclusion criteria to participate. Four were unable to complete the MRI portion of the study, resulting in the final sample of twenty-four OAs. Recruitment for the OAs took place through community outreach as well as online recruitment engines including craigslist.org and BuildClinical.com. Participants were included if they 1) were at least 50 years old, 2) had normal hearing, and 3) passed MRI screening. Participants were excluded if they 1) changed medications within 6 weeks of screening; 2) had a history of psychotic or schizophrenic episodes, major neurological diagnosis, or other medical condition that might impair cognition; 3) had a history of chemotherapy within the past 10 years; or 4) experienced serious physical trauma or were diagnosed with a serious chronic health condition requiring medical treatment and monitoring within 3 months of screening. Participants were compensated for their time through either payment or course credit. This study was approved by the Northeastern University Institutional Review Board.

For Hypothesis 3, predictions pertaining to music listening were tested using the same sample as above, while predictions not specifically relating to task incorporated additional resting state data as described below. These additional data included 22 YA and OA participants from the music listening study not included in the other analyses, and these participants met the same inclusion criteria listed above. The other N = 58 participants’ data were taken from previous studies (Belden et al., 2020; Przysinda et al., 2017), resulting in a total sample of N = 128. In this study, adults aged 18-39 participated in return for monetary compensation or course credit. Participants were recruited from Wesleyan University, Hartt School of Music, Northeastern University, Berklee College of Music, and the local community in the Greater Boston area, and included selection for participants with and without musical training experience.

### Behavioral Measures

Participants reported demographic data, including age, gender, race, and household income level. Participants also completed the Barcelona Music Reward Questionnaire (Mas-Herrero et al., 2014), a twenty-question survey in which participants rated their overall levels of musical reward through subscales of emotional evocation, mood regulation, sensory motor, social reward, and music seeking. Each subscale had four questions pertaining to it, and each question was rated on a 5-point Likert scale, including 3 reverse-scored items.

### MRI Data Acquisition

Images were acquired using a Siemens Magnetom 3T MR scanner with a 64-channel head coil at Northeastern University. For task fMRI data, continuous acquisition was used for 1440 volumes with a fast TR of 475 ms, for a total acquisition time of 11.4 minutes. Forty-eight axial slices (slice thickness = 3 mm, anterior to posterior, z volume = 14.4 mm) were acquired as echo-planar imaging (EPI) functional volumes covering the whole brain (TR = 475 ms, TE = 30 ms, flip angle = 60°, FOV = 240mm, voxel size = 3 × 3 × 3 mm3).

The resting state scan followed the same parameters and included 947 continuous scans, for a total scan length of approximately 7 and a half minutes. T1 images were also acquired using a MPRAGE sequence, with one T1 image acquired every 2400 ms, for a total task time of approximately 7 minutes. Sagittal slices (0.8 mm thick, anterior to posterior) were acquired covering the whole brain (TR = 2400 ms, TE = 2.55 ms, flip angle = 8°, FOV= 256, voxel size = 0.8 × 0.8 × 0.8 mm3) (as described in Quinci et al., 2022).

### Functional Music Listening Task

The fMRI task consisted of 24 trials altogether. In each trial, participants were first presented with a musical stimulus (lasting 20 seconds), then they were given the task of rating how familiar they found the music to be (familiarity rating lasted 2 seconds), and how much they liked the music (liking rating also lasted 2 seconds). Musical stimuli for the MRI task consisted of 24 different audio excerpts, each of which was 20 seconds in duration. Each audio stimulus was from one of the following three categories: participant self-selected music (6/24 stimuli), researcher-selected music including well-known excerpts spanning multiple musical genres (10/24 stimuli) and novel music spanning multiple genres (8/24 stimuli). Stimuli were presented in a randomized order, and participants made ratings of familiarity and liking on the scales of 1 to 4: for familiarity: 1=very unfamiliar, 2=unfamiliar, 3=familiar, 4=very familiar; for liking: 1=hate, 2=neutral, 3=like, 4=love. Participants made these ratings by pressing a corresponding button on a button-box (Cambridge Research Systems) inside the scanner. Participants wore MR-compatible over-the-ear headphones (Cambridge Research Systems) over musician-grade silicone ear plugs during MRI data acquisition. The spatial mapping between buttons and the numerical categories of ratings were counterbalanced between subjects to reduce any systematic association between familiarity or liking and the motor activity resulting from making responses.

### MRI Preprocessing

Task and resting state fMRI data were preprocessed using the CONN Toolbox (Whitfield-Gabrieli & Nieto-Castanon, 2012). Preprocessing steps included functional realignment and unwarp, functional centering, functional slice time correction, functional outlier detection using the artifact detection tool, functional direct segmentation and normalization to MNI template, structural centering, structural segmentation and normalization to MNI template, and functional smoothing to an 8mm gaussian kernel (Friston et al., 1995). Denoising steps included white matter and cerebrospinal fluid confound correction (Behzadi et al., 2007), motion correction, global signal regression, and bandpass filtering to 0.008-0.09 Hz.

### Graphical Feature Extraction

First-level, preprocessed and denoised task and resting state fMRI timeseries data were extracted for each ROI of the CONN Toolbox default atlas. Timeseries data were then correlated across ROIs to form two connectivity matrices for each subject, one for music listening and one for rest. Given expected differences in overall connection strengths between our two samples, we used an absolute threshold for our connectivity matrices rather than a proportional threshold. For statistical testing, we used an absolute threshold of r = 0.3, which showed global connectivity patterns characteristic of the overall relationship between groups and conditions. Graph theoretical measures of modularity, global efficiency, clustering coefficients, degrees, and betweenness centrality were extracted using the Brain Connectivity Toolbox (BCT) (Rubinov & Sporns, 2010) for each subject and each condition. Modularity was defined as the total number of modules identified for each subject and each condition, and global measures of clustering coefficients, degrees, and betweenness centrality were defined as the average of these measures across all ROIs.

### Analysis Plan

To address Hypothesis 1, a group-by-task MANCOVA was performed with global measures of modularity, Eglob, CC, BC, and degrees as dependent variables. This was modeled as a repeated-measures general linear model, with task as the repeated measure IV, group as the main between-subjects IV, and covariates of BMRQ, Gender, and within-group age, defined as subject age minus the average age of their group.

To address Hypothesis 2, an ROI-by-group-by-task MANCOVA was performed with local measures of CC, BC, and degrees in mPFC (Medial Frontal Cortex from CONN atlas) and PCC (Posterior Cingulate from CONN atlas) as dependent variables. This was modeled using the same parameters as the global measures MANCOVA, with the only difference being inclusion of ROI as an additional repeated-measure IV.

To address Hypothesis 3, additional MANCOVA analyses were performed for resting state data only in the larger sample of 128. These were defined as a Multivariate GLM in SPSS, and used the same dependent variables reported in the Global Measures MANCOVA. Participants were tertile split according to their scores on the BMRQ, forming groups of musical anhedonics (ANH, lowest third), hedonics (H, middle third), and hyper-hedonics (HH, highest third) to function as the main between-subjects IV. Linear covariates of age and gender were also included. The global-measures MANCOVA was performed for the full CONN atlas, and then repeated using only the subset of ROIs identified as the auditory and reward systems by previous work in our group (Wang et al., 2020), for which network measures were extracted separately using the same methodology reported above.

Results from all MANCOVA models will be presented as follows: Summary statistics for all between-subject factors, within-subject factors, covariates, and interactions between these measures will be reported first. Then, for any factors showing significance at the uncorrected p < 0.05 level, follow-up analyses will identify which network measures are driving the effect. These results are considered significant only if p < 0.05 after performing Benjamini-Hochberg correction for all comparisons included at this level of analysis, using a False Detection Rate of Q = 0.05. The p-values in this report are uncorrected, but all effects presented did survive correction unless otherwise noted. Summary tables for these models are included in the supplementary materials at https://osf.io/2je98/.

## Results

### Older Adults Show Less Modular Network Structures in Favor of More Globally Connected Graphs

The OA group showed higher degrees, higher clustering, higher betweenness centrality, and lower modularity than the YA group, with between-group effects being stronger during music listening than during rest, as shown in Figures 1 and 2. Overall, the global measures group-by-task MANCOVA model was significant for main effects of group (F(5,39) = 4.971, p = 0.001, partial η^2^ = 0.389) and task (F(5,39) = 2.511, p = 0.046, partial η^2^ = 0.243). Significant interactions were also observed between task and group (F(5,39) = 6.025, p < 0.001, partial η^2^ = 0.436), and between task and within-group age (F(5,39) = 4.643, p = 0.001, partial η^2^ = 0.373).

**Figure 1:**
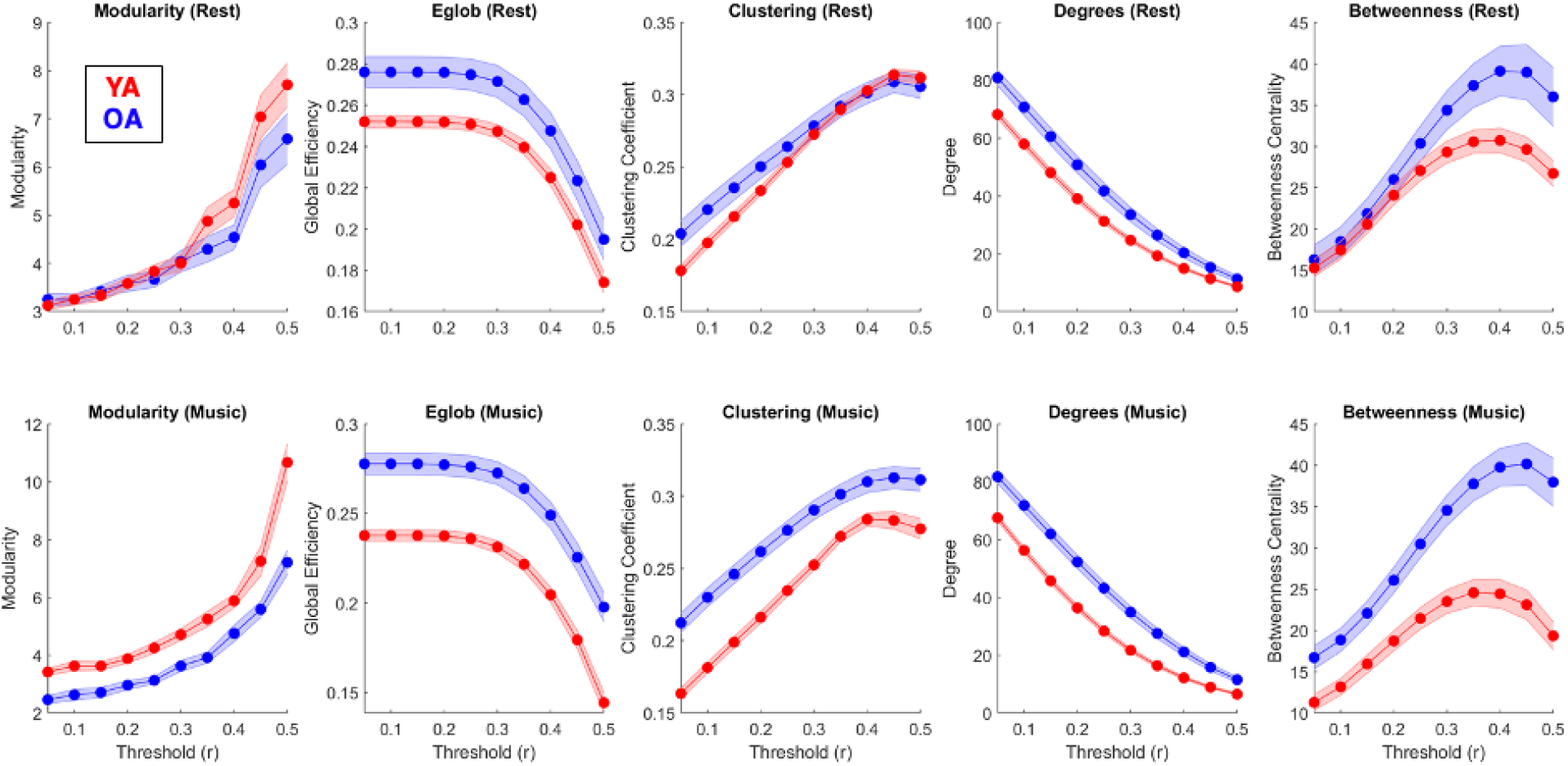
Age x Task Cross-Threshold GT Measures. Global measures of modularity, global efficiency, clustering coefficients, degrees, and betweenness centrality in YAs (red) and OAs (blue) during rest (top) and music listening (bottom). Lines depict the mean value of each measure across all nodes and all subjects across a range of correlation thresholds, and the shaded region represents the between-subject standard error for each group.

**Figure 2:**
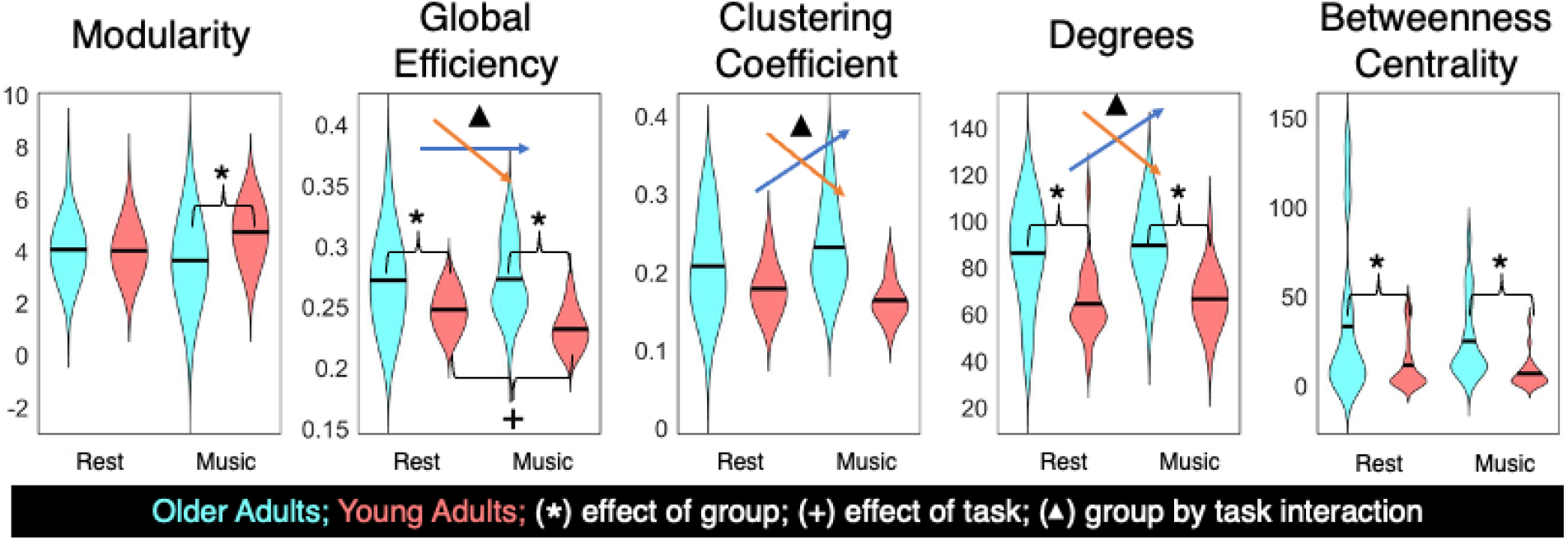
Age x Task Global Measures MANCOVA. Summary of effects comparing YAs (red) and OAs (blue) between tasks in global measures of modularity, global efficiency, clustering coefficient, degree, and betweenness centrality in medial prefrontal cortex and posterior cingulate cortex. All effects are significant at the p < 0.05, Benjamini-Hochberg corrected level.

Significant effects of this model are summarized in Figure 2. Main effects of group were characterized by differences in modularity (F(5,39) = 9.719, p = 0.003, partial η^2^ = 0.184), global efficiency (F(5,39) = 9.202, p = 0.004, partial η^2^ = 0.176), degrees (F(5,39) = 10.814, p = 0.002, partial η^2^ = 0.201), and betweenness centrality (F(5,39) = 7.279, p = 0.010, partial η^2^ = 0.145). While modularity was higher in YAs as compared to OAs, measures of Eglob, degrees, and BC were all higher in the OA sample.

### Music Listening Reinforces Age-Specific Connectivity Profiles

A main effect of task was observed for global efficiency, such that Eglob was higher during rest than during music listening (F(5,39) = 5.213, p = 0.027, partial η^2^ = 0.108). This effect is further explained by an interaction between group and task (F(5,39) = 6.875, p = 0.012, partial η^2^ = 0.138), in which Eglob decreased during music listening in YAs but not OAs. Similar interaction effects were also present for CC (F(5,39) = 25.682, p < 0.001, partial η^2^ = 0.374), and degrees (F(5,39) = 5.638, p = 0.022, partial η^2^ = 0.116), although these were characterized by increases during music listening in OAs in addition to the decreases observed in YAs.

Task by within-sample age interactions were also observed for Eglob (F(5,39) = 5.617, p < 0.022, partial η^2^ = 0.116), CC (F(5,39) = 9.118, p < 0.004, partial η^2^ = 0.175), and degrees (F(5,39) = 10.180, p < 0.003, partial η^2^ = 0.191), such that for each age group, there was a positive effect of age on all these measures during rest, and a negative effect during music listening.

**Table 1:** Highest Ranked Regions for Local Measures. Regions showing the highest group-averaged CC, degrees, and BC during rest and music listening conditions. Legend: anterior cingulate cortex (AC); anterior superior temporal gyrus (aSTG); cerebellum 4/5 (Cereb45); cerebellum 6 (Cereb6); central opercular cortex (CO); frontal orbital cortex (FOrb); Heschl’s gyrus (HG); insular cortex (IC); intracalcarine cortex (ICC); inferior frontal gyrus pars opercularis (IFG oper); lingual gyrus (LG); middle frontal gyrus (MidFG); occipital fusiform gyrus (OFusG); occipital pole (OP); paracingulate gyrus (PaCiG); posterior cingulate cortex (PC); posterior middle temporal gyrus (pMTG); parietal operculum (PO); postcentral gyrus (PostCG); planum polare (PP); parietal parahippocampal cortex (pPaHC); precentral gyrus (PreCG); parietal superior temporal gyrus (pSTG); planum temporale (PT); supracalcarine cortex (SCC); supplementary motor area (SMA); superior parietal lobule (SPL); temporo-occipital inferior temporal gyrus (toITG); temporo-occipital middle temporal gyrus (toMTG); temporal pole (TP).

| Rank | OA Clustering |  | YA Clustering |  | OA Degrees |  | YA Degrees |  | OA Betweenness |  | YA Betweenness |  |
| --- | --- | --- | --- | --- | --- | --- | --- | --- | --- | --- | --- | --- |
| Rest |  |  |  |  |  |  |  |  |  |  |  |  |
| 1 | CO r | 0.363 | SCC r | 0.411 | IC l | 52.667 | IC l | 38.917 | LG r | 127.583 | LG r | 107.083 |
| 2 | Cuneal r | 0.359 | ICC r | 0.406 | HG l | 51.625 | IC r | 38.750 | AC | 122.000 | Cereb45 l | 96.583 |
| 3 | SCC l | 0.358 | ICC l | 0.404 | IC r | 50.583 | PP l | 35.667 | IC r | 116.333 | Cereb6 l | 85.917 |
| 4 | CO l | 0.357 | Cuneal r | 0.399 | HG r | 50.417 | PP r | 35.583 | IC l | 112.000 | IC r | 84.917 |
| 5 | ICC r | 0.354 | LG r | 0.397 | AC | 50.250 | HG l | 34.917 | LG l | 110.333 | IC l | 77.417 |
| 6 | PO r | 0.351 | SCC l | 0.395 | PT r | 48.958 | TP r | 34.708 | PT l | 96.667 | LG l | 77.083 |
| 7 | ICC l | 0.350 | LG l | 0.394 | PT l | 48.542 | Cereb45 l | 34.125 | Cereb6 l | 87.333 | PP r | 73.250 |
| 8 | SCC r | 0.347 | Cuneal l | 0.392 | PP r | 48.333 | HG r | 33.542 | PreCG l | 84.833 | PT l | 70.833 |
| 9 | SMA r | 0.344 | OFusG l | 0.368 | FOrb r | 47.458 | IFG oper r | 32.500 | PT r | 80.250 | PP l | 64.583 |
| 10 | PreCG r | 0.340 | OP r | 0.366 | PP l | 47.375 | Cereb6 l | 32.375 | PaCiG r | 79.583 | Cereb6 r | 60.167 |
| Music Listening |  |  |  |  |  |  |  |  |  |  |  |  |
| 1 | aSTG l | 0.365 | SCC r | 0.439 | IC l | 52.625 | pMTG r | 34.625 | LG l | 140.333 | pMTG r | 95.583 |
| 2 | pSTG r | 0.363 | ICC l | 0.416 | IC r | 52.000 | TP r | 33.125 | LG r | 139.417 | PP l | 82.500 |
| 3 | pSTG l | 0.362 | SCC l | 0.404 | Precuneous | 51.542 | FOrb r | 32.917 | Cereb6 r | 107.750 | PT l | 78.000 |
| 4 | aSTG r | 0.350 | ICC r | 0.392 | toMTG l | 50.958 | pMTG l | 32.708 | pMTG r | 103.833 | pSTG r | 67.583 |
| 5 | PP l | 0.340 | Cuneal l | 0.390 | PC | 50.667 | PP l | 30.708 | IC r | 101.500 | PP r | 64.750 |
| 6 | Cuneal r | 0.339 | Cuneal r | 0.390 | FOrb r | 49.833 | FOrb l | 30.625 | PT l | 94.083 | PT r | 64.750 |
| 7 | SCC r | 0.338 | LG r | 0.365 | MidFG l | 48.500 | IC l | 30.542 | Cereb6 l | 90.250 | LG l | 59.750 |
| 8 | ICC r | 0.333 | LG l | 0.365 | pMTG l | 48.333 | Cereb45 l | 30.458 | IC l | 88.000 | aSTG r | 58.417 |
| 9 | PostCG r | 0.332 | aSTG r | 0.360 | AC | 48.167 | TP l | 30.250 | PT r | 86.000 | Cereb45 l | 58.083 |
| 10 | SPL r | 0.331 | PT l | 0.345 | toITG l | 47.542 | pPaHC l | 29.667 | PP l | 84.500 | IC l | 57.917 |

### Regional Connectivity Profiles Differ by Age and Task

While measures of CC, degrees, and BC averaged across all ROIs were higher in OAs than YAs, only the latter two of these reached significance in the global measures MANCOVA. One explanation for this is the broader range of CC in YAs between ROIs: For example, across both conditions, the 5 ROIs with the highest average CC across all YA participants were higher in this measure than the five ROIs with the highest average CC in the OA group (Table 3). The opposite is true for degrees and BC, for which the top 5 in OA are consistently higher than the top 5 in YA.

Figure 3 shows network statistics across the brain regions for each age group and for each task. Visual inspection of the figure shows that the regions with consistently high CC in YA are primarily visual processing regions, including bilateral supracalcarine and intracalcarine cortices. The auditory regions in the superior temporal lobe showed higher clustering during music listening as compared to rest in both groups, as expected. Furthermore, this clustering increase in auditory regions during the music-listening task was especially pronounced for the OA group, such that the bilateral anterior and posterior superior temporal gyrus (STG) became the regions with the highest average CC across the entire brain during music listening, surpassing even the visual processing regions which are otherwise typically highly clustered. In contrast, for the YA group, the right anterior STG was the only auditory network region to be in the top 10 highest CCs during music listening. Other notable regions within the auditory network were the right posterior MTG, which showed the highest degrees and BC during music listening in YAs, and bilateral Heschl’s Gyrus (HG), which showed the highest degrees in the OAs during rest. Meanwhile, bilateral Insular Cortex (IC) showed consistently high degrees during both rest and music-listening in both groups.

**Figure 3:**
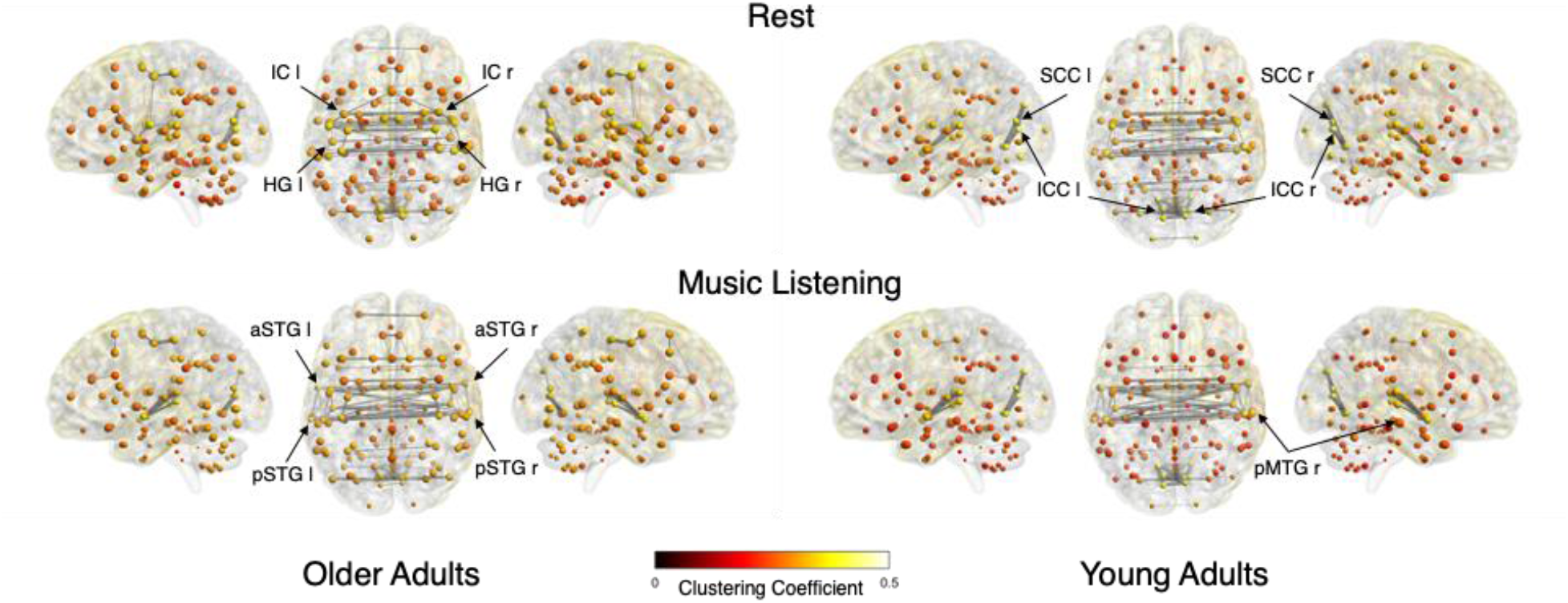
Connectome Visualizations. Visualizations of young (right) and OA (left) connectomes during rest (top) and music listening (bottom). In all cases, node size is scaled by degree, and node color is scaled by clustering coefficient (see colorbar). All nodes were mapped, but only the top 10% of edges are shown. Labeled nodes include those showing highest degree and/or clustering coefficient in the corresponding group and condition.

### PCC is More Central than mPFC in Older Adults

Overall, the local measures MANCOVA model was significant for main effects of group (F(3,41) = 6.645, p < 0.001, partial η^2^ = 0.327) and ROI (F(3,41) = 10.072, p<0.001, partial η^2^ = 0.424). There was a significant two-way interaction between ROI and task (F(3,41) = 5.722, p < 0.002, partial η^2^ = 0.295), as well as significant three-way ROI by task by group (F(3,41)=2.928 p<0.045, partial η^2^ = 0.176) and ROI by task by within-sample-age (F(3,41)=3.566, p<0.022, partial η^2^ = 0.207) interactions.

Specific effects of this model are summarized in Figure 4. A main effect of group was observed in degrees (F(3,41) = 7.144, p < 0.011, partial η^2^ = 0.142), with this measure being higher in OAs than YAs. A main effect of ROI was also observed in both degrees (F(3,41) = 15.061, p < 0.001, partial η^2^ = 0.259) and BC (F(3,41) = 31.16, p < 0.001, partial η^2^ = 0.420), with these measures being higher in PCC than mPFC. Degrees were also specifically higher during music listening in PCC, leading to a ROI by task interaction (F(3,41) = 9.74, p = 0.003, partial η^2^ = 0.185). These effects were primarily driven by the OA group; however the main effect of group in BC (F(3,41) = 5.461, p = 0.024, partial η^2^ = 0.113) and ROI by group by task interaction in degrees (F(3,41) = 4.604, p = 0.032, partial η^2^ = 0.097) did not survive Benjamini-Hochberg correction.

**Figure 4:**
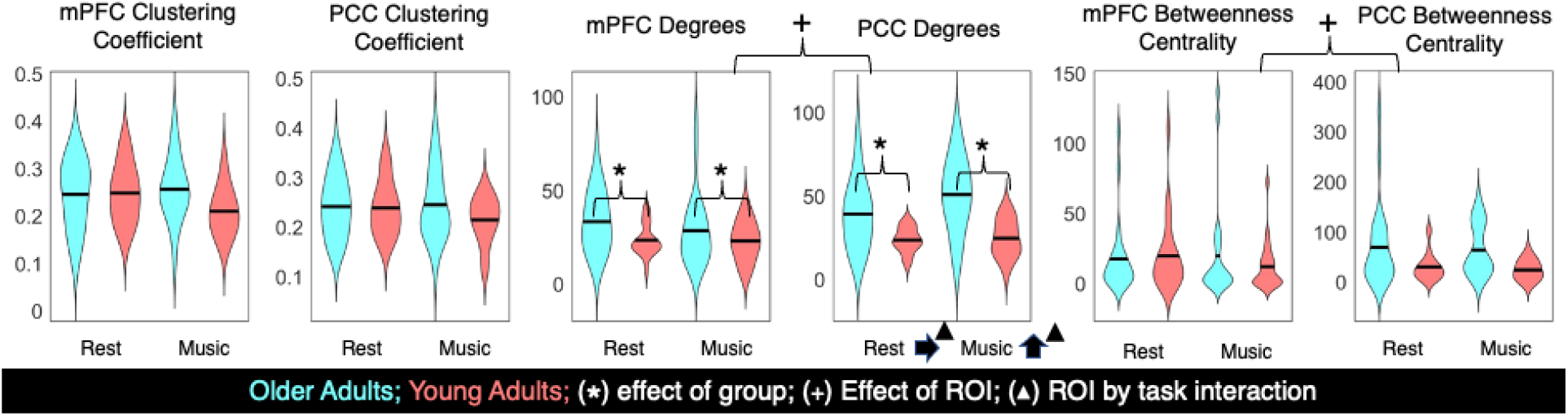
Age x Task x ROI Local Measures MANCOVA. Summary of effects comparing YAs (red) and OAs (blue) between tasks in measures of clustering coefficient, degree, and betweenness centrality in medial prefrontal cortex and posterior cingulate cortex. All effects are significant at the p < 0.05, Benjamini-Hochberg corrected level.

### Differences in Musical Reward Do Not Modulate Graph Structure at Rest

The overall model was not significant for any main effects or interactions relating to BMRQ for both the whole brain parcellation and auditory-reward network specific parcellation. In addition, BMRQ was included as a linear covariate in the previously reported Global and Local Measures MANCOVA models, and was also found not to be significant.

## Discussion

We analyzed the functional brain topology in YAs and OAs, both at rest and during music listening, and related graph theory metrics to between-group and between-task differences. We found significant age-related reorganization of brain networks, and age-by-task interactions for both whole-brain and DMN specifically, thus supporting our first two hypotheses. We did not find support for individual differences in musical reward affecting the topology of whole-brain or default mode networks. Taken together, results represent a novel characterization of the modulatory effects of music on age-related differences in the functional connectome.

Our first prediction was that modularity would decrease with age, but that music listening would recover modular function specifically in OAs, leading to a group by task interaction. While we did observe a main effect of group on modularity favoring YAs, this difference was primarily observed in the music listening, and not the rest condition. This suggests that the more modular connectivity profile observed in YAs is more pronounced in the presence of music. It is worth noting that this may be a more general effect of task-based fMRI, and at this time more testing is needed. In addition, we observed further main effects of group suggesting that OAs have a more globally efficient connectome than YAs, while previous findings have not shown this difference (Geerligs et al., 2015). However, this previous report was limited to resting state data, while here the music condition may have improved the visibility of latent age-related connectivity differences.

Second, we expected that OAs would show a shift in connectivity from mPFC to PCC, resulting in increased BC and decreased CC in PCC relative to YAs, and the opposite effect in mPFC. No significant effects were observed in CC, but a main effect of ROI on BC was observed. While the effect of group on BC did not survive correction, it is clear that the higher BC observed in PCC relative to mPFC was primarily driven by the OA sample, following the predicted pattern. Unlike the predicted pattern, the CC and BC of mPFC did not differ between groups. Degrees were overall higher in OA than YA, reflective of patterns observed in the global measures MANCOVA. In addition, there was an ROI-by-task interaction indicating that degrees were higher during music listening than rest in PCC, but not mPFC. Similar to the marginal effect of group in BC, this was primarily driven by differences observed in the OA group, such that PCC becomes even more highly connected in OAs when listening to music. This suggests that while PCC is more hublike in OAs relative to YAs, it may not be compensating for a dysregulated mPFC. Further studies should take a deeper look into the specific role of PCC in aging.

The findings support different organizing principles of the functional connectome in young and OAs: The YA connectome is characterized by a tighter, more modular structure, led by densely connected sensory regions. In contrast, the OA connectome is characterized by a high level of global connectivity, leading to increased efficiency at the cost of modularity. Strikingly, in the presence of music, these differences in connectivity profile become even more pronounced. The YA connectome becomes more modular at the cost of global efficiency, degrees, and CC, while OAs increase their CC and degrees without sacrificing their high global efficiency. What remains a question is how these connections relate to cognitive aging measures. For example, while network dysregulation is known to be related to cognitive decline (Buckner et al., 2005), work has also shown some areas of improved cognitive function in OAs (Veríssimo et al., 2022). At this time, it is unclear whether the differential connectivity patterns between age groups reflect connectivity deficits related to cognitive decline, or instead a change in cognitive strategy that may be beneficial in some ways. For this reason, we suggest future studies attempt to link these observed differences in connectomic structure to known behavioral differences between young and OAs.

Finally, we did not find any support for our third hypothesis, that differences in musical reward will lead to more highly connected networks, especially among auditory and reward regions. BMRQ was not shown to be significant in any of our models, either in a tertile split or when used as a linear covariate in the group-by-task analyses. This suggests that musical reward sensitivity may not have a straightforward bearing on the structure of the functional connectome, even when responding to music, and that the graphical structure of auditory and reward systems is largely similar across musical anhedonics, hedonics, and hyperhedonics.

While we have previously seen that listening to music that is well-liked, and especially music self-selected by the participant, leads to increased functional connectivity between auditory and reward regions (Belden et al., 2023), the present result suggests that the overall connective structure of auditory and reward systems is not necessarily dependent on trait-level musical reward, at least as it is captured using the self-report measure of the BMRQ. While previous work has shown differences in connectivity between auditory and reward regions varying with self-reported musical reward in both brain structure and brain function (Martínez-Molina et al., 2016; Sachs et al., 2016; Salimpoor et al., 2013), it is possible that those with higher trait-level musical reward do in fact show increased connectivity between auditory and reward regions relative to their less musical-reward sensitive peers, but that these differences do not result in a significant reorganization of the overall connective structure of these systems, as captured using graph theoretical tools.

Taken together, these findings further our understanding of the effects of aging on the functional connectome, and how rewarding stimuli such as music can modulate these effects. Over the course of aging, there seems to be a shift in connectomic structure, from a more modular profile in young adulthood to one that is instead defined by heightened global efficiency for task-related processes in older adulthood. Engagement with music listening is a task that transiently bolsters these differences, regardless of the listener’s personal degree of general musical reward experience.

Still, these findings bring with them a large number of follow-up questions. For example, it is currently unclear why the PCC is so much more richly connected in OAs, and whether or not it should be treated as a target for future MBI’s. Above all, as this work was primarily focused on effects of group and task on brain network organization, more work is needed to draw conclusions about how these different brain connectivity profiles may be linked to reward learning behavior, and to cognitive function more generally. Drawing the link between these observed functional network dynamics and behavioral changes across the lifespan remains a critical step for the development of effective music-based interventions for OAs with or at risk for AD and other forms of cognitive decline.

## Acknowledgements

This work was supported by funding from National Institutes of Health (NIH R01AG078376, NIH R21AG075232) and National Science Foundation (NSF-BCS 2240330, and NSF-CAREER 1945436) to PL.

## References

Aalto, S., Brück, A., Laine, M., Någren, K., & Rinne, J. O. (2005). Frontal and Temporal Dopamine Release during Working Memory and Attention Tasks in Healthy Humans: A Positron Emission Tomography Study Using the High-Affinity Dopamine D2 Receptor Ligand [11C]FLB 457. Journal of Neuroscience, 25(10), 2471–2477. https://doi.org/10.1523/JNEUROSCI.2097-04.2005

Ai, M., Loui, P., Morris, T. P., Chaddock-Heyman, L., Hillman, C. H., McAuley, E., & Kramer, A. F. (2022). Musical Experience Relates to Insula-Based Functional Connectivity in Older Adults. Brain Sciences, 12(11), Article 11. https://doi.org/10.3390/brainsci12111577

Alluri, V., Toiviainen, P., Burunat, I., Kliuchko, M., Vuust, P., & Brattico, E. (2017). Connectivity patterns during music listening: Evidence for action-based processing in musicians. Human Brain Mapping, 38(6), 2955–2970. https://doi.org/10.1002/hbm.23565

Bäckman, L., Lindenberger, U., Li, S.-C., & Nyberg, L. (2010). Linking cognitive aging to alterations in dopamine neurotransmitter functioning: Recent data and future avenues. Neuroscience & Biobehavioral Reviews, 34(5), 670–677. https://doi.org/10.1016/j.neubiorev.2009.12.008

Badgaiyan, R. D., Fischman, A. J., & Alpert, N. M. (2007). Striatal dopamine release in sequential learning. NeuroImage, 38(3), 549–556. https://doi.org/10.1016/j.neuroimage.2007.07.052

Behzadi, Y., Restom, K., Liau, J., & Liu, T. T. (2007). A component based noise correction method (CompCor) for BOLD and perfusion based fMRI. NeuroImage, 37(1), 90–101. https://doi.org/10.1016/j.neuroimage.2007.04.042

Belden, A., Quinci, M. A., Geddes, M., Donovan, N. J., Hanser, S. B., & Loui, P. (2023). Functional Organization of Auditory and Reward Systems in Aging. Journal of Cognitive Neuroscience, 1–23. https://doi.org/10.1162/jocn_a_02028

Belden, A., Zeng, T., Przysinda, E., Anteraper, S. A., Whitfield-Gabrieli, S., & Loui, P. (2020). Improvising at rest: Differentiating jazz and classical music training with resting state functional connectivity. NeuroImage, 207, 116384. https://doi.org/10.1016/j.neuroimage.2019.116384

Buckner, R. L., Snyder, A. Z., Shannon, B. J., LaRossa, G., Sachs, R., Fotenos, A. F., Sheline, Y. I., Klunk, W. E., Mathis, C. A., Morris, J. C., & Mintun, M. A. (2005). Molecular, Structural, and Functional Characterization of Alzheimer’s Disease: Evidence for a Relationship between Default Activity, Amyloid, and Memory. Journal of Neuroscience, 25(34), 7709–7717. https://doi.org/10.1523/JNEUROSCI.2177-05.2005

Chen, W. G., Iversen, J. R., Kao, M. H., Loui, P., Patel, A. D., Zatorre, R. J., & Edwards, E. (2022). Music and Brain Circuitry: Strategies for Strengthening Evidence-Based Research for Music-Based Interventions. Journal of Neuroscience, 42(45), 8498–8507. https://doi.org/10.1523/JNEUROSCI.1135-22.2022

Chowdhury, R., Guitart-Masip, M., Lambert, C., Dayan, P., Huys, Q., Düzel, E., & Dolan, R. J. (2013). Dopamine restores reward prediction errors in old age. Nature Neuroscience, 16(5), Article 5. https://doi.org/10.1038/nn.3364

Eppinger, B., Nystrom, L. E., & Cohen, J. D. (2012). Reduced Sensitivity to Immediate Reward during Decision-Making in Older than Younger Adults. PLOS ONE, 7(5), e36953. https://doi.org/10.1371/journal.pone.0036953

Erixon-Lindroth, N., Farde, L., Robins Wahlin, T.-B., Sovago, J., Halldin, C., & Bäckman, L. (2005). The role of the striatal dopamine transporter in cognitive aging. Psychiatry Research: Neuroimaging, 138(1), 1–12. https://doi.org/10.1016/j.pscychresns.2004.09.005

Fang, R., Ye, S., Huangfu, J., & Calimag, D. P. (2017). Music therapy is a potential intervention for cognition of Alzheimer’s Disease: A mini-review. Translational Neurodegeneration, 6(1), 2. https://doi.org/10.1186/s40035-017-0073-9

Farahani, F. V., Karwowski, W., & Lighthall, N. R. (2019). Application of Graph Theory for Identifying Connectivity Patterns in Human Brain Networks: A Systematic Review. Frontiers in Neuroscience, 13. https://www.frontiersin.org/articles/10.3389/fnins.2019.00585

Friston, Karl. J., Ashburner, J., Frith, C. D., Poline, J.-B., Heather, J. D., & Frackowiak, R. S. J. (1995). Spatial registration and normalization of images. Human Brain Mapping, 3(3), 165–189. https://doi.org/10.1002/hbm.460030303

Gaviola, M. A., Inder, K. J., Dilworth, S., Holliday, E. G., & Higgins, I. (2020). Impact of individualised music listening intervention on persons with dementia: A systematic review of randomised controlled trials. Australasian Journal on Ageing, 39(1), 10–20. https://doi.org/10.1111/ajag.12642

Geddes, M. R., Mattfeld, A. T., Angeles, C. de los, Keshavan, A., & Gabrieli, J. D. E. (2018). Human aging reduces the neurobehavioral influence of motivation on episodic memory. NeuroImage, 171, 296–310. https://doi.org/10.1016/j.neuroimage.2017.12.053

Geerligs, L., Renken, R. J., Saliasi, E., Maurits, N. M., & Lorist, M. M. (2015). A Brain-Wide Study of Age-Related Changes in Functional Connectivity. Cerebral Cortex, 25(7), 1987–1999. https://doi.org/10.1093/cercor/bhu012

Global Council on Brain Health. (2020). Music on Our Minds: The Rich Potential of Music to Promote Brain Health and Mental Well-Being. Global Council on Brain Health. https://doi.org/10.26419/pia.00103.001

Grady, C., Sarraf, S., Saverino, C., & Campbell, K. (2016). Age differences in the functional interactions among the default, frontoparietal control, and dorsal attention networks. Neurobiology of Aging, 41, 159–172. https://doi.org/10.1016/j.neurobiolaging.2016.02.020

Hafkemeijer, A., van der Grond, J., & Rombouts, S. A. R. B. (2012). Imaging the default mode network in aging and dementia. Biochimica et Biophysica Acta (BBA) - Molecular Basis of Disease, 1822(3), 431–441. https://doi.org/10.1016/j.bbadis.2011.07.008

Karmonik, C., Brandt, A., Anderson, J. R., Brooks, F., Lytle, J., Silverman, E., & Frazier, J. T. (2016). Music Listening Modulates Functional Connectivity and Information Flow in the Human Brain. Brain Connectivity, 6(8), 632–641. https://doi.org/10.1089/brain.2016.0428

Klostermann, E. C., Braskie, M. N., Landau, S. M., O’Neil, J. P., & Jagust, W. J. (2012). Dopamine and frontostriatal networks in cognitive aging. Neurobiology of Aging, 33(3), 623.e15-623.e24. https://doi.org/10.1016/j.neurobiolaging.2011.03.002

Knutson, B., Fong, G. W., Bennett, S. M., Adams, C. M., & Hommer, D. (2003). A region of mesial prefrontal cortex tracks monetarily rewarding outcomes: Characterization with rapid event-related fMRI. NeuroImage, 18(2), 263–272. https://doi.org/10.1016/S1053-8119(02)00057-5

Koepp, M. J., Gunn, R. N., Lawrence, A. D., Cunningham, V. J., Dagher, A., Jones, T., Brooks, D. J., Bench, C. J., & Grasby, P. M. (1998). Evidence for striatal dopamine release during a video game. Nature, 393(6682), Article 6682. https://doi.org/10.1038/30498

Landau, S. M., Lal, R., O’Neil, J. P., Baker, S., & Jagust, W. J. (2009). Striatal Dopamine and Working Memory. Cerebral Cortex, 19(2), 445–454. https://doi.org/10.1093/cercor/bhn095

Laukka, P. (2007). Uses of music and psychological well-being among the elderly. Journal of Happiness Studies, 8(2), 215–241. https://doi.org/10.1007/s10902-006-9024-3

Leggieri, M., Thaut, M. H., Fornazzari, L., Schweizer, T. A., Barfett, J., Munoz, D. G., & Fischer, C. E. (2019). Music Intervention Approaches for Alzheimer’s Disease: A Review of the Literature. Frontiers in Neuroscience, 13. https://www.frontiersin.org/articles/10.3389/fnins.2019.00132

Li, S.-C., & Rieckmann, A. (2014). Neuromodulation and aging: Implications of aging neuronal gain control on cognition. Current Opinion in Neurobiology, 29, 148–158. https://doi.org/10.1016/j.conb.2014.07.009

MacDonald, S. W. S., Li, S.-C., & Bäckman, L. (2009). Neural underpinnings of within-person variability in cognitive functioning. Psychology and Aging, 24(4), 792–808. https://doi.org/10.1037/a0017798

Martínez-Molina, N., Mas-Herrero, E., Rodríguez-Fornells, A., Zatorre, R. J., & Marco-Pallarés, J. (2016). Neural correlates of specific musical anhedonia. Proceedings of the National Academy of Sciences, 113(46), E7337–E7345. https://doi.org/10.1073/pnas.1611211113

Mas-Herrero, E., Marco-Pallares, J., Lorenzo-Seva, U., Zatorre, R. J., & Rodriguez-Fornells, A. (2014). Barcelona Music Reward Questionnaire [dataset]. https://doi.org/10.1037/t31533-000

Mather, M. (2016). The Affective Neuroscience of Aging. Annual Review of Psychology, 67(1), 213–238. https://doi.org/10.1146/annurev-psych-122414-033540

Monchi, O., Hyun Ko, J., & Strafella, A. P. (2006). Striatal dopamine release during performance of executive functions: A [11C] raclopride PET study. NeuroImage, 33(3), 907–912. https://doi.org/10.1016/j.neuroimage.2006.06.058

O’Doherty, J., Winston, J., Critchley, H., Perrett, D., Burt, D. M., & Dolan, R. J. (2003). Beauty in a smile: The role of medial orbitofrontal cortex in facial attractiveness. Neuropsychologia, 41(2), 147–155. https://doi.org/10.1016/S0028-3932(02)00145-8

Onoda, K., Ishihara, M., & Yamaguchi, S. (2012). Decreased Functional Connectivity by Aging Is Associated with Cognitive Decline. Journal of Cognitive Neuroscience, 24(11), 2186–2198. https://doi.org/10.1162/jocn_a_00269

Onoda, K., & Yamaguchi, S. (2013). Small-worldness and modularity of the resting-state functional brain network decrease with aging. Neuroscience Letters, 556, 104–108. https://doi.org/10.1016/j.neulet.2013.10.023

Palomar-García, M.-Á., Zatorre, R. J., Ventura-Campos, N., Bueichekú, E., & Ávila, C. (2016). Modulation of Functional Connectivity in Auditory–Motor Networks in Musicians Compared with Nonmusicians. Cerebral Cortex, bhw120. https://doi.org/10.1093/cercor/bhw120

Przysinda, E., Zeng, T., Maves, K., Arkin, C., & Loui, P. (2017). Jazz musicians reveal role of expectancy in human creativity. Brain and Cognition, 119, 45–53. https://doi.org/10.1016/j.bandc.2017.09.008

Quinci, M. A., Belden, A., Goutama, V., Gong, D., Hanser, S., Donovan, N. J., Geddes, M., & Loui, P. (2022). Longitudinal changes in auditory and reward systems following receptive music-based intervention in older adults. Scientific Reports, 12(1), Article 1. https://doi.org/10.1038/s41598-022-15687-5

Rubinov, M., & Sporns, O. (2010). Complex network measures of brain connectivity: Uses and interpretations. NeuroImage, 52(3), 1059–1069. https://doi.org/10.1016/j.neuroimage.2009.10.003

Sachs, M. E., Ellis, R. J., Schlaug, G., & Loui, P. (2016). Brain connectivity reflects human aesthetic responses to music. Social Cognitive and Affective Neuroscience, 11(6), 884–891. https://doi.org/10.1093/scan/nsw009

Sala-Llonch, R., Bartrés-Faz, D., & Junqué, C. (2015). Reorganization of brain networks in aging: A review of functional connectivity studies. Frontiers in Psychology, 6. https://www.frontiersin.org/articles/10.3389/fpsyg.2015.00663

Salimpoor, V. N., van den Bosch, I., Kovacevic, N., McIntosh, A. R., Dagher, A., & Zatorre, R. J. (2013). Interactions Between the Nucleus Accumbens and Auditory Cortices Predict Music Reward Value. Science, 340(6129), 216–219. https://doi.org/10.1126/science.1231059

Samanez-Larkin, G. R., Worthy, D. A., Mata, R., McClure, S. M., & Knutson, B. (2014). Adult age differences in frontostriatal representation of prediction error but not reward outcome. Cognitive, Affective, & Behavioral Neuroscience, 14(2), 672–682. https://doi.org/10.3758/s13415-014-0297-4

Schott, B. H., Minuzzi, L., Krebs, R. M., Elmenhorst, D., Lang, M., Winz, O. H., Seidenbecher, C. I., Coenen, H. H., Heinze, H.-J., Zilles, K., Düzel, E., & Bauer, A. (2008). Mesolimbic Functional Magnetic Resonance Imaging Activations during Reward Anticipation Correlate with Reward-Related Ventral Striatal Dopamine Release. Journal of Neuroscience, 28(52), 14311–14319. https://doi.org/10.1523/JNEUROSCI.2058-08.2008

Schultz, W., Dayan, P., & Montague, P. R. (1997). A Neural Substrate of Prediction and Reward. Science, 275(5306), 1593–1599. https://doi.org/10.1126/science.275.5306.1593

Sheline, Y. I., Raichle, M. E., Snyder, A. Z., Morris, J. C., Head, D., Wang, S., & Mintun, M. A. (2010). Amyloid Plaques Disrupt Resting State Default Mode Network Connectivity in Cognitively Normal Elderly. Biological Psychiatry, 67(6), 584–587. https://doi.org/10.1016/j.biopsych.2009.08.024

Tomasi, D., & Volkow, N. D. (2012). Aging and functional brain networks. Molecular Psychiatry, 17(5), Article 5. https://doi.org/10.1038/mp.2011.81

Ueda, T., Suzukamo, Y., Sato, M., & Izumi, S.-I. (2013). Effects of music therapy on behavioral and psychological symptoms of dementia: A systematic review and meta-analysis. Ageing Research Reviews, 12(2), 628–641. https://doi.org/10.1016/j.arr.2013.02.003

Veríssimo, J., Verhaeghen, P., Goldman, N., Weinstein, M., & Ullman, M. T. (2022). Evidence that ageing yields improvements as well as declines across attention and executive functions. Nature Human Behaviour, 6(1), Article 1. https://doi.org/10.1038/s41562-021-01169-7

Wang, D., Belden, A., Hanser, S. B., Geddes, M. R., & Loui, P. (2020). Resting-State Connectivity of Auditory and Reward Systems in Alzheimer’s Disease and Mild Cognitive Impairment. Frontiers in Human Neuroscience, 14. https://www.frontiersin.org/articles/10.3389/fnhum.2020.00280

Whitfield-Gabrieli, S., & Nieto-Castanon, A. (2012). Conn: A Functional Connectivity Toolbox for Correlated and Anticorrelated Brain Networks. Brain Connectivity, 2(3), 125–141. https://doi.org/10.1089/brain.2012.0073

Yee, D. M., Adams, S., Beck, A., & Braver, T. S. (2019). Age-Related Differences in Motivational Integration and Cognitive Control. Cognitive, Affective, & Behavioral Neuroscience, 19(3), 692–714. https://doi.org/10.3758/s13415-019-00713-3

Zhang, L., Wang, X., Alain, C., & Du, Y. (2023). Successful aging of musicians: Preservation of sensorimotor regions aids audiovisual speech-in-noise perception. Science Advances, 9(17), eadg7056. https://doi.org/10.1126/sciadv.adg7056

